# In silico neuritogenesis model underpins mechanical interactions with extracellular matrix as determinants of persistent axonal growth in stiffer microenvironments

**DOI:** 10.64898/2026.03.13.708543

**Authors:** Mathar Kravikass, Lars Bischof, Kristina Karandasheva, Federica Furlanetto, Pritha Dolai, Sven Falk, Marisa Karow, Katja Kobow, Ben Fabry, Vasily Zaburdaev

## Abstract

It has been broadly recognized that the crosstalk between cells and their extracellular matrix (ECM) is crucial for the proper function of biological tissues. Relatively recently the role of ECM came in focus in the context of neuronal development and regeneration, where the effects of the ECM mechanics on the migration of neurons and neurite growth are still incompletely understood. Here we present an in silico twin framework for neurite growth focusing on its biophysical interactions with the ECM. This coarsegrained model accounts for viscoelastic liquid- and solid-like ECMs and neurite growth by ECM-mediated traction forces. Resulting growth trajectories can be rationalized based on the theory of random walks and polymer physics. To critically assess model’s predictive power, we performed experiments on neurites of hippocampal rat neurons growing in 3D collagen gels and observed a more persistent axon outgrowth in denser matricies. The model fully recapitulated the effect, thereby underpinning the central role of mechanical interactions with ECM as guiding principle of axonal growth. We argue that a combination our model with optical microscopy may provide an is silico twin helping to disentangle the contributions of “passive” physics from more complex effects of chemical queues or an apparent mechanosensing.

## 1 Introduction

During the early stages of brain development, neurons differentiate and extend protrusions: axons and dendrites, collectively known as neurites, which actively explore their environment in search of synaptic partners [1]. The resulting growth patterns shape the final structure of the neuronal network, which, in turn, is closely linked to overall neural function [2, 3]. Consequently, developmental pathways must be robust and reproducible to ensure reliable brain tissue function across individuals.

Mechanistically, neurite motility is governed by the growth cone, a dynamic structure at the neurite’s tip tasked with environmental sensing and forward extension. Growth cone mobility arises from a combination of substrate adhesion and cytoskeletal remodeling, with local actin dynamics generating protrusive forces and adhesive contacts stabilizing forward motion [4, 5, 6]. The neurite shaft itself is supported by bundled microtubules, rigid cylindrical protein assemblies arranged along neurite’s length [4, 5]. As the neurite elongates, microtubules polymerize within the shaft and exert forces against the growth cone, creating a mechanical feedback that regulates extension [5].

In addition to intrinsic mechanisms, it is well established that the extracellular matrix (ECM) plays a crucial role in shaping neural development at all stages [7]. During neurite extension, the ECM provides both guidance cues and a physical substrate for neuronal growth. While chemical signals are known to influence growth trajectories [1, 5, 8], mechanical properties such as stiffness, adhesion strength, and topology also significantly affect neurite development. This influence extends well beyond the stages of development, where it still greatly impacts plasticity-related functions such as memory retention [9]. Identifying these mechanical cues is challenging because their effects often overlap and intertwine with chemical signals. An example of such a relation is how the mechanosensing ion channel Piezo1 suppresses transthyretin transport protein expression on stiffer substrates, resulting in a delay in voltage-gated ion channel activity, action potentials, and synapse formation, all essential for enabling neuronal communication [10].

The generic biophysical argument for the importance of mechanics in growth is that the primary mode of motility ultimately depends on the forces exerted on or by neurites. The growing neurite can therefore be described as a mechanically regulated system in which both internal tension within the neurite and growth cone extension play key roles [5]. Consequently, theoretical modeling of neurites and their growth has become increasingly important for describing these physical interactions. Proposed models vary in both complexity and in their interpretation of which mechanisms are most critical to the process [11]. For example, a recent multiscale model demonstrates that distinct mechanical processes operate at different levels: at the molecular scale, substrate stiffness governs axonal growth cone interactions, while at the cellular scale, axon mechanics can be described using a morphoelastic filament framework [12]. Another recent model instead focuses on substrate mechanics arising from adhesive interactions between neurites and the ECM, as determined by the geometry of the matrix itself [13].

In addition to advances in numerical modeling, recent *in vivo* and *in vitro* techniques show promise in deciphering mechanical processes in neural development. Brain organoid models enable targeted studies of human neurons derived from pluripotent stem cells [14], while animal models continue to provide a crucial foundation for investigating interactions occurring during growth [15]. With growing recognition of the ECM as a key regulator of neurite growth, numerous studies have focused on characterizing these relationships [7, 16], employing ECMs derived from animal tissues, as well as commercially available matrices such as MatriGel [16]. Additionally, artificial ECMs made out of hydrogels such as collagen offer reproducible, controlled environments in which morphogenetic events of neuronal growth can be examined in isolation, focusing on substrate interactions [17].

While a broader range of techniques such as atomic force microscopy (AFM) [1, 8], traction force microscopy and novel imaging techniques like Interferometric Scattering Microscopy (iSCAT) [18] could provide rich data output for individual neurite tracking, the effects of the mechanical interactions with the ECM still remain much less understood than chemical signaling pathways [5].

Therefore, here we start with proposing a model which will act as an *in silico* twin framework that aims to disentangle biophysical underpinnings in a variety of the experimental settings. This work builds upon the original study dissecting the effect of genetic mutations on developmental defects by linking it to the biomechanics of neurite growth [19]. Here, our generalized framework is designed to be integrated seamlessly with a broad range of experimental systems and can accommodate a variety of ECM substrates and geometries. Aiming at high throughput and parameter scanning, highly-detailed models of neurites [11, 12] become computationally impractical and limit analytical tractability of the results. Instead, we reference a simplified growth model in which the neurite is represented as a polymer-like chain with tension-based interactions between evenly spaced joints [5]. This abstraction provides two key advantages: it facilitates the incorporation of complex environmental influences and allows us to leverage established theoretical frameworks from polymer physics.

We use this model to demonstrate how the mechanical crosstalk of neurite and ECM shapes growth trajectories in a systematic dependence on a small number of biologically tractable parameters. Dynamics and morphology of neurite growth can be quantified and rationalized by using the theoretical concepts of random walks and polymer physics.

Next, to critically test the predictive power of the model, we performed experiments quantifying axonal outgrowth of hippocampal rat neurons in 3D collagen gels with varying concentrations. We discovered that in gels with higher collagen content the neurites grow with higher persistence, while the speed of the outgrowth is not strongly affected. We could successfully use the model to quantitatively recapitulate how ECMs with higher volume fraction of the polymer and stiffness scaling with ECMs polymer content results in a more persistent outgrowth of axons. Thus, the “passive” physics of axon-ECM interactions is sufficient to describe the effect without invoking the “active” mechanism of mechanosensing. We argue that our image- and simulations-based framework can be expanded to other, more complex processes of brain tissue development and regeneration. In combination with other imaging modalities, it will enable to quantify and disentangle the effects of biophysical interactions on the neurite outgrowth and thus help to capture the effects due to biochemical signaling and active biological regulation. We next proceed to describe the model setup.

## 2 Numerical Model

Our model was originally motivated by the experiments performed *in vitro* on brain organoid systems where the neurite growth was quantified in dependence on the genetic mutations corresponding to a specific neurodevelopmental disorder (see [19] for details). In brief, organoids derived from the respective pluripotent stem cell lines and their isogenic control lines were cut in thin slices, put on the supporting Matrigel so that the growth of multiple isolated neurites could be observed as a function of time by optical microscopy in an effectively two-dimensional setting (see **Figure 1a**). This experiment provided a good basis on the relevant number of neurites to be investigated, their morphology and time scales. The ultimate goal is to use the comparison between the model and experiment under varying conditions to pinpoint and test the responsible microscopic changes in neurite-ECM interactions [19]. Therefore, motivated by this organoid data (Figure 1a) we propose the following coarse grained model of the neurito-genesis. For the sake of simplicity, we provide the description for the two-dimensional scenario, while the same model with minimal adjustments is also applicable to the three-dimensional case.

**Figure 1.**
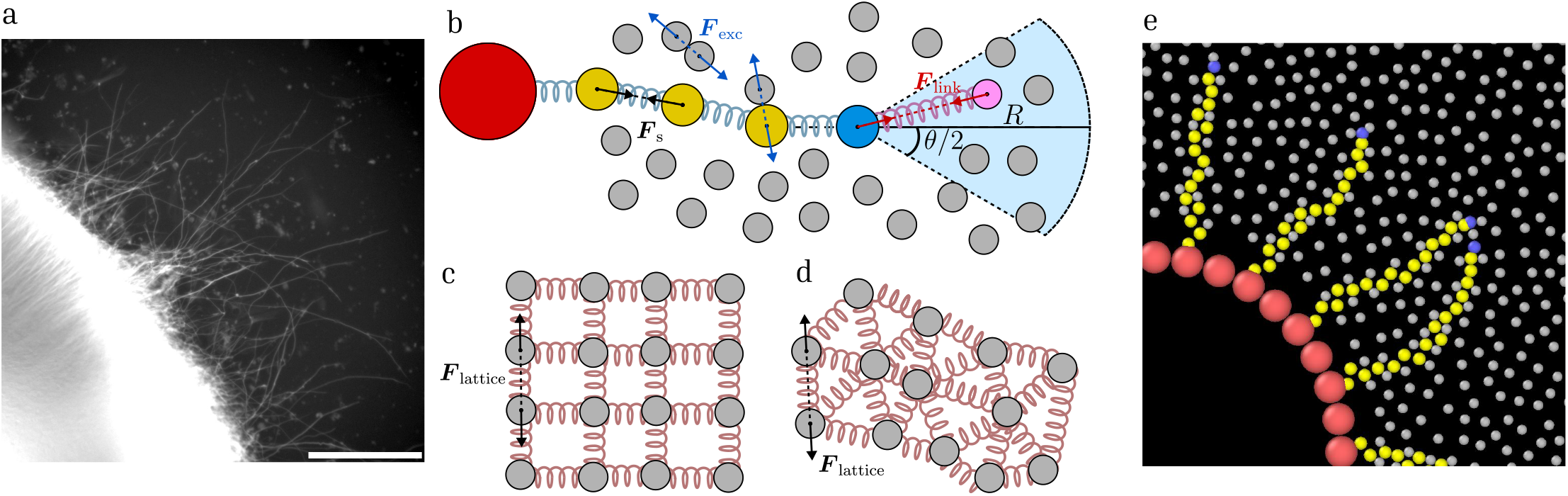
a) Live imaging snapshot of neurite outgrowth from an organoid slice. Scale bar = 200*µm*. Image shown was exposure corrected to emphasize the protruding neurites. See [19] for further details. b) Schematic of the agent-based model. The neurites are represented by a chain of (yellow) beads connected by springs protruding out of the neurite cell body (red) and growing in the extracellular space occupied by (grey) beads representing the extracellular matrix (ECM). The growth cone bead of the neurite (blue) can establish a temporary link with a matrix particle (magenta), randomly chosen within a certain arc angle *θ* and range *R* in front of the neurite. The linked matrix particle then connects to the leading bead by a harmonic spring. Meanwhile, as the neurites advance through the matrix, its geometry is slowly deformed by repulsive excluded volume interactions. b) ECM particles connected by springs in a square lattice configuration. d) ECM particles with random initial positions, connected by Delaunay triangulation. e) Simulation snapshot of the model. See Supplementary Video 1.

Neuron cell bodies are modeled as immobile circles with radius *R*_neuron_. Protruding from the neuron body, the neurites are modeled by a chain of beads (Figure 1b) connected by harmonic springs. This simplification draws on the basic assumption that neurites are maintained by tension forces dependent on the dynamics between the neuron body and the growth cone driving the extension of the neurite [5]. The beads representing the neurite body are all of equal radius *R*_neurite_ and mobility *µ*_neurite_. The springs connecting the beads have coefficient *κ*_neurite_ and equilibrium length *l*_neurite_, leading to the force between each two neighboring beads *i* and *i* + 1:

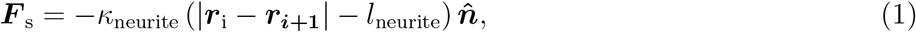

where ***r***_i_ is the position vector of the *i*-th bead and 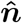 is the unit vector in the direction towards the next bead. The very first bead of the growing chain corresponds to the growth cone of the neurite. Growth of the neurite occurs by the elongation of the chain element that connects the first bead to the rest of the chain. When this chain element length reaches the equilibrium length *l*_neurite_, the bead becomes a part of the neurite body, and the new bead is introduced in the same position to drive the next step of the growth. We will distinguish between two growth modes: ECM-independent and ECM-dependent, relying on the adhesion and traction force generation. During the growth, the spring constant of the first spring is set to a lower value *κ*_cone_ to allow for elongation by traction forces. When the first bead is introduced, the equilibrium length of the spring is set to 0. In the ECM-independent growth mode, which mimics slow intrinsic neurite elongation, this equilibrium length grows at constant rate *α*. To introduce the ECM-dependent growth, we first briefly describe how the matrix is modeled.

The ECM is represented as a set of circular particles (disks) of radius *R*_ECM_. This representation is dictated by numerical efficiency and yet it also reflects the recent experimental efforts where artificial ECMs were recapitulated by densely packed gel beads with diameter in the order of 15*µm* [20]. The beads are placed randomly with no overlap at a given number density (area fraction *ϕ*_ECM_). This type of ECM would correspond to the case of a viscoelastic fluid-like material (Figure 1b). By connecting the ECM beads by harmonic springs with beads either arranged in a regular lattice-like configuration (Figure 1c) or randomly (Figure 1d) we can also describe a viscoelastic solid-like environment. The mobility of the ECM particles *µ*_ECM_ is chosen with consideration of Stoke’s law.

Active interactions of the growth cone with the surrounding ECM enable neurite extension. We assume that this interaction can be represented by a traction force between the ECM particles and the forefront bead of the chain. To set up the link, a random ECM particle is chosen within the maximum range *R* and inside of the arc with opening angle *θ*. The symmetry axis of the arc is along the direction connecting the forefront and the previous bead, and thus regulates the persistence of growth direction. The forefront bead and the selected ECM bead get connected by a harmonic spring with zero equilibrium length and spring constant *κ*_link_ which represents the traction force acting on both beads:

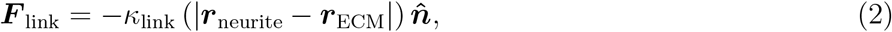

where 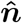 is the direction from the linked matrix particle to the forefront bead acting on the neurite and the same with the opposite sign acting on the ECM particle. This traction force causes both the forefront and the ECM particle to move towards each other, thus extending the chain link leading to the forefront bead. Once this length reaches the target link length, the new chain bead is introduced and forefront particle update happens as described before. The neurite-ECM link is formed with a fixed persistence time of *t*_link_. After the link to the ECM breaks, a new random ECM particle is chosen and the process continues. Movement of the ECM beads and those representing neurite lead to the interactions between beads when they overlap, which we model as excluded volume harmonic force with the spring constant *ϵ*:

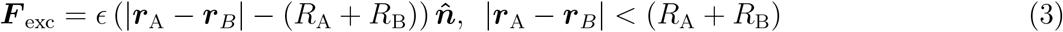

where 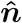 is the direction connecting the centers of beads A and B respectively. For simplicity and mimicking relevant experimental setups we will not consider excluded volume interaction for the neurite beads and thus will allow them to intersect each other.

Finally, in the overedamped limit (where we neglect inertia forces compared to spring, excluded volume, and friction forces, which is justified at this microscopic scale) the displacement in each simulation step *dt* of the growth cone is given by:

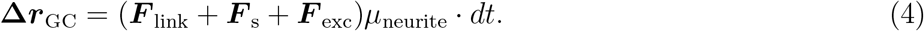

Meanwhile, for the neurite beads:

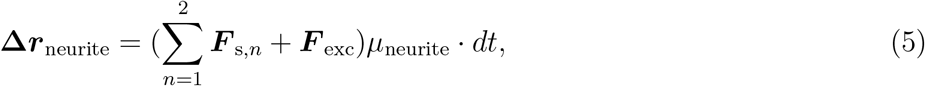

Where ***F***_s,*n*_ is the force exerted from neighbor *n* from either side of the bead. In the solid-like configuration of the ECM, the harmonic springs connecting the matrix beads exert the following force on each bead *i* given its neighbor ensemble *j* = 0, 1…*N* :

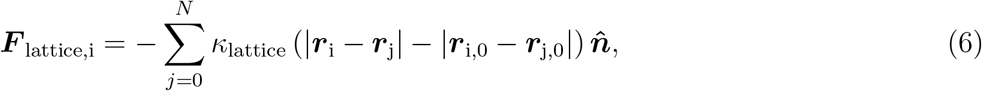

where *κ*_lattice_ is the lattice spring coefficient, ***r***_i,0_ and ***r***_j,0_ are the initial positions of beads *i* and *j* respectively, and 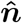 is the direction vector between the two beads. Finally, the displacement of a bound ECM particle is given by:

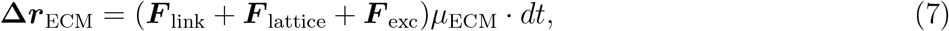

where for the case of a non-linked particle or a non-connected ECM, ***F***_link_ and ***F***_lattice_ will be 0, respectively. In practice, to focus on the phenotypes of neurite outgrowth, neuron bodies are placed in the center of the simulation box (Figure 1e), as to both allow each neurite equal space to grow, and to more closely represent experimental configurations in the organoid system (Figure 1a).

To facilitate a dimensionless formulation, all simulation parameters are expressed relative to a set of reference scales rather than carrying intrinsic physical units. These reference scales define the quantities that are taken to be unity in the normalized system, and all other parameters are scaled accordingly by their dimensional relationship to them. As such, scaling the simulation parameters to match experimental observations can be achieved by scaling the reference scales with known quantities. The spatial scale is set by the characteristic diameter of a neurite, which is chosen as the fundamental unit of length and fixed to 2*R*_neurite_ = 1.0. Force scales are normalized using a reference spring coefficient *κ* associated with growth cone–adhesion interactions. Specifically, *κ* is defined such that a displacement of one unit length between a growth cone and an adhesion site produces a traction force of magnitude 1.0. Since this traction force is experimentally measurable (typically ~ nN [21]), it provides a physically meaningful force scaling. Finally, the simulation time unit is defined by the time required for a particle of radius *R* to move its own diameter under the action of a unit force. This time scale is dependant on the mobility parameter by *t*_0_ = 2*R/µF*, from which we set *µ* = 1.0. In this normalization, the reference spring constant therefore satisfies *κ* = 1, and all other spring parameters are expressed relative to this value.

## 3 Results

In this section, we first provide detailed investigation of the theoretical model and its behavior in a broad range of the model parameters together with the possible theoretical interpretation by means of polymer physics theory and the model of random walks. We next report our experimental findings on the 3D *in vitro* system of rat hippocampal neurons growing their axons in the collagen gel ECM. We show that the increasing of ECM polymer density makes the axonal growth more persistent. We then show how the model quantitatively recapitulates these results.

### Liquid-like extracellular matrix

First, we considered a liquid-like extracellular matrix in which the ECM particles can move freely in space under the action of forces generated by growing neurites. We use this setup to investigate how the model parameters determine phenotypes of neurite growth and introduce an analytical theory rooted in polymer physics and random walks to quantify this growth.

We identified the growth sector adhesion range parameters *θ* and *R* to have a significant impact on neurite growth. In particular, variations in *θ* directly affect the directionality of growth. Thus, we first investigate the role of *θ* by performing simulations of changing angles from 10° to 60°. The neurons are placed in a circle at the middle of the simulation box, such that 13 actively growing neurites are simulated in each run. The area fraction of the ECM particles in the simulation box was chosen to be *ϕ*_ECM_ = 0.8, where the radius of each ECM particle was chosen as *R*_ECM_ = 0.25, half of the neurite radius. The rest of the growth sector parameters were kept at the following constants: *R* = 20, *t*_link_ = 0.1, *κ*_link_ = 1.0. For a full list of the parameters, see Table S1. Finally, the box size of the simulation was set as 400, chosen to ensure neurites do not reach the edges of the simulation space.

To characterize the growth trajectories, we track the *x, y* coordinates of the forefront bead as a function of time. The trajectory of the forefront bead in the broad range of parameters is quantitatively very similar to the final outline of the neurite (Supplementary Figure S1). The tracking occurs by sampling a set number of frames out of the simulation, which we chose as 1000 out of a total of 20000 simulation steps. While our main goal in this sampling was to reduce complexity without damaging the results, this too can be modified to better fit the measurement rate of an experiment. Due to this subsampling, we now use the sampling step *dt* instead of the simulation time steps for the definitions below. We start with quantifying the mean squared displacement (MSD) of the forefront bead, which we calculate for a set of *N* trajectories:

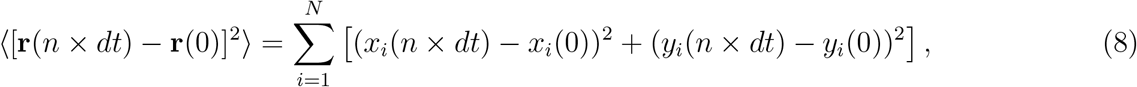

Where *n* is the discretized time step, and *x*_*i*_(*t*), *y*_*i*_(*t*) are the *x, y* axis displacement of the *i*-th trajectory at time *t* = *n* × *dt*, respectively. We observe that the MSD decreases with increasing *θ*, indicating reduced directional persistence of the growth (Figure 2a). The MSD exhibits two regimes as clearly visible in the log-log scale plot. The initial scaling with time is ∝ *t*^2^ reflecting the persistent, ballistic-like motion, while after some characteristic time it switches to the diffusive scaling ∝ *t*. This characteristic transition time is lower for larger *θ*.

**Figure 2.**
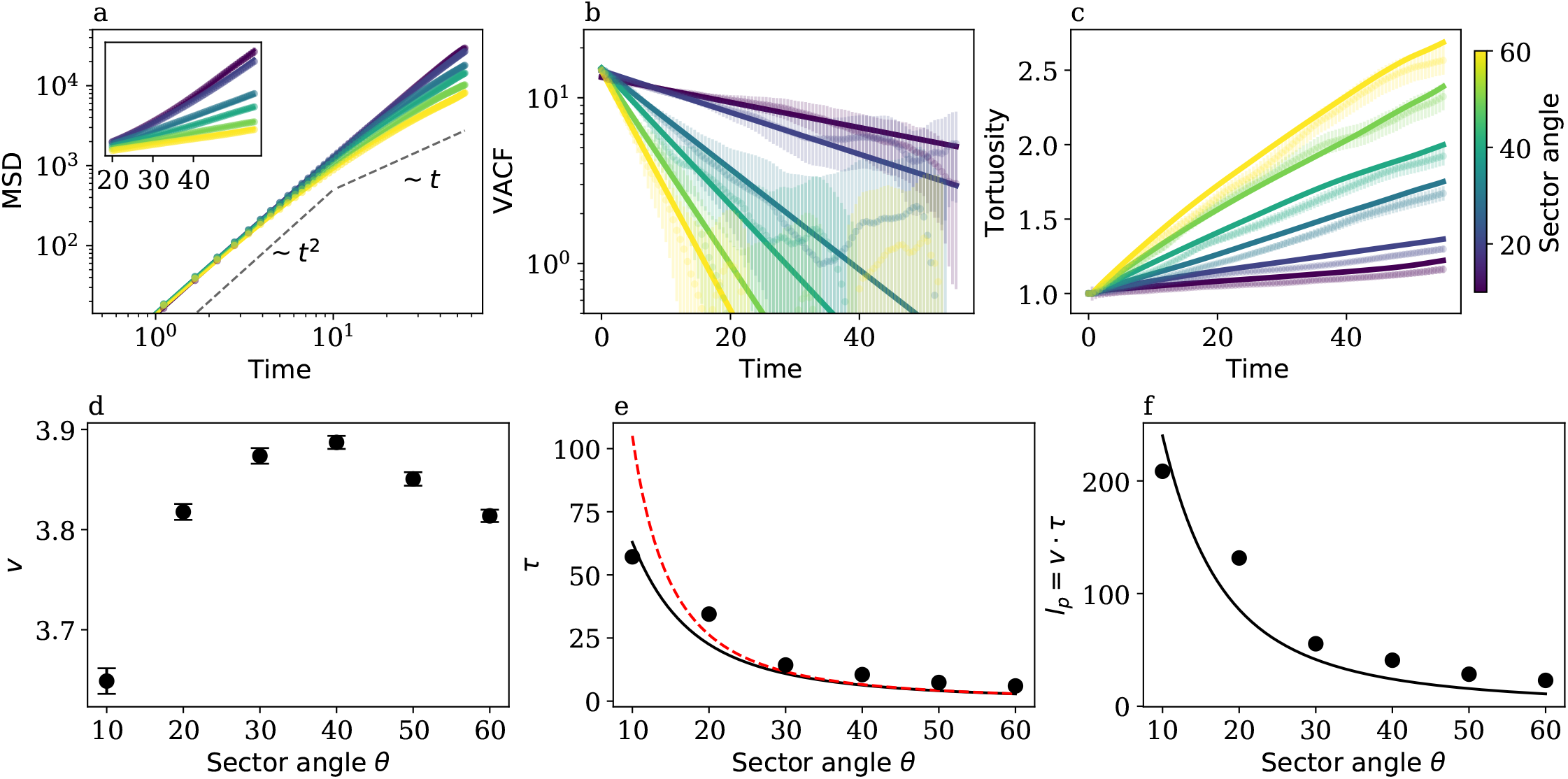
Simulation results averaged over 130 trajectories per sector angle *θ*, using the parameters described in the main text. Colors correspond to the simulated *θ*, as indicated by the color bar. Error bars represent the standard error. a) Mean squared displacement (MSD) as a function of time, shown on a double logarithmic scale for each *θ*. Solid colored lines represent fitted theoretical MSD (Equation 13). Dashed black lines indicate the change in MSD scaling from an initial ballistic regime (~ *t*^2^) to a diffusive regime (~ *t*^1^). The characteristic crossover time shifts toward *t* = 0 as *θ* increases. Inset: MSD over time in linear for *t* ≥ 20. b) Velocity autocorrelation function (VACF) as a function of time on semi-logarithmic scale. The logarithm of the VACF scales linearly with log *t*, in agreement with theoretical predictions (Equation (11)). Solid colored lines represent the fitted theoretical VACF. c) Tortuosity as a function of time for different *θ* values. Solid lines indicate fitting to the theoretical prediction of 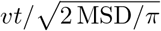. d) Root mean squared (RMS) velocity *v* extracted from the simulations and used in the model fitting. e) Fitted persistence time *τ* as a function of sector angle. The red line shows the run-and-tumble theoretical prediction (Equation 12), while the solid black line shows the modified version (Equation 14). f) Persistence length *l*_*p*_ as a function of sector angle. The solid line represents the theoretical prediction of (*l*_*p*_) assuming a constant velocity equal to the average over all simulated angles.

We can also compute the velocity autocorrelation function (VACF), which quantifies how fast the growth velocity at time *t* loses its correlation with the initial value at time *t*_0_, defined as:

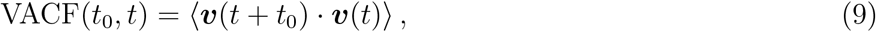

Where ***v***(*t*) is the velocity vector of the trajectories at time step *t*. In order to reduce the effects of microscopic fluctuations we defined the velocity by the increments of the coordinates divided by the increments of time, with a delay of 20 *dt*, where *dt* is the sampled timestep. The results are not sensitive to this delay within a broad range (Supplementary Figure S2). The VACFs show an exponential decay with faster velocity decorrelation at larger sector angles (Figure 2b). We also calculated tortuosity (trajectory contour length divided by the end-to-end distance) for each trajectory. Intuitively, higher sector angles lead to more “curved” trajectories which is also reflected in higher tortuosity values (Figure 2c).

We can get further insights into the observed behavior by using simple models of statistical physics. Specifically, in the field of polymer physics, when describing configurations of a polymer in space, and in the closely related theory of random walks investigating statistics of the trajectories as a function of time. A well-known random walk model is that of a run-and-tumble [22, 23, 24, 25]. Here, a particle follows a trajectory consisting of straight runs alternated by stochastic (instantaneous) reorientation events (tumbles) which change the direction of motion. Motion happens with constant speed *v* while the durations of runs are often considered to be exponentially distributed. Turning angle, defined as the angle between two consecutive runs, is random, following a certain distribution, determines the persistence of movements and leads to a correlated random walk model. In the simplest version, the turning angles are chosen to be random with respect to the previous direction and given by ± *θ*_RT_. When *θ*_RT_ = 0 the particle always moves along the straight line, while *θ*_RT_ = *π/*2 corresponds to an uncorrelated random walk (see below). A closely related model of freely rotating chain [26] is known for the description of polymer configurations in space. Here, the monomer length is equivalent to the length of a single run, and the bending angle between the two monomers is equivalent to the turning angle. One key difference is that the monomers have a fixed constant length, unlike the typically used exponentially distributed lengths of the runs. Both models successfully describe the phenomenon of directional correlation in the orientation of runs/monomers and a transition to a fully random-walk-like (diffusive) behavior after a large enough number of runs or monomers in the chain. The total temporal duration of the trajectory in the run-and-tumble corresponds to the length of the polymer chain in the polymer model. One notable advantage of the run-and-tumble model is its ability to provide an exact analytical description of both the mean squared displacement (MSD) and the velocity autocorrelation function (VACF) as functions of time (due to the exponential statistics of runs). For this reason, we adopt this phenomenological framework here. We next briefly summarize these results and argue how the phenomenological results of the run-and-tumble model can be related to the microscopic parameters of our neurite growth model. We assume that the run times are exponentially distributed with a mean *τ*_run_:

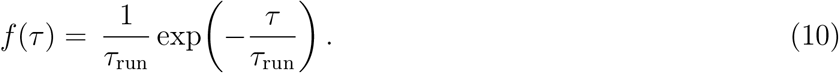

From here, the formulation of the VACF is well known and has a characteristic exponential decay [24]:

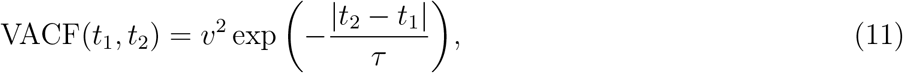

where the characteristic decay time (persistence time) *τ* is determined by *τ*_run_ and *θ*_RT_ via:

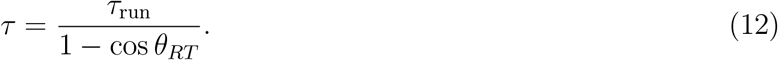

The MSD can be then derived via the Green-Kubo relation as a double temporal integral of the VACF leading to:

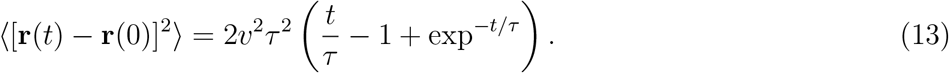

This resulting expression is the well-known formula for the MSD of the Ornstein-Uhlenbeck process [27]. We now can use this low-parametric formula to quantify the neurite growth trajectories from our simulations. The average growth velocity *v* used for the model was calculated by the root mean squared velocity of the simulated neurites. We see that velocities vary by less than 10% across angles, indicating no significant correlation between sector angle and speed (Figure 2d). We next fitted the VACF and MSD for the different sector angles to the model prediction (Equation 11) with the persistence time *τ* as a fit parameter. For the MSD, the model replicates the switch from ballistic (~ *t*^2^) to diffusive (~ *t*^1^) scalings observed in the simulation data accurately (Figure 2a). It can be additionally seen that the characteristic time of the switch decreases as the sector angle is increased. The same shift in persistence scaling can be seen in the VACF plot, where higher sector angles exhibit a steeper downward linear slope in semi-logarithmic scale, aligning with lower *τ* values (Figure 2b). The persistence time *τ* is then compared to the theoretical prediction (Equation 10), where *τ*_run_ was taken as the link persistence time *t*_link_.

In qualitative agreement with the analytical result, higher turning angles during growth result in trajectories with a lower persistence length. In the ideal run-and-tumble process, the correlation time diverges as the angle *θ* approaches 0. In our simulation model, however, elongations are not perfectly straight as the bead trajectory is affected by the collisions with the ECM particles, which can be quantified by an independent process of rotational diffusion. This effect is then incorporated in the run-and-tumble model and predicts the effective correlation time to be a geometric mean of decorrelation due to diffusion, *τ*_RD_ and due to run-and-tumble, *τ*, [22]:

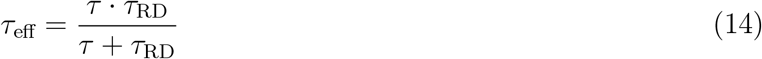

By using *τ*_RD_ as a fit parameter while *τ*_run_ remains a constant *t*_link_, we arrive to an approximation of the correlation time from our simulations which aligns better for the lower angles (Figure 2e).

Finally, we can also calculate the persistence length of the trajectory as *l*_*p*_ = *v τ* (Figure 2f). As *v* has only a minor dependence on the sector angle, a theoretical prediction for *l*_*p*_ was constructed by multiply-ing the model of *τ* (Equation 14) by *v* averaged over all angles.

The results on the correlation time also translate to the values of the effective diffusion constant when considering the longer time *t* ≫ *τ*, when the MSD becomes:

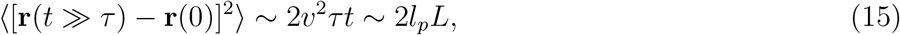

where *L* is the contour length, given by *L* = *vt*. This result corresponds to the mean squared end-to-end distance of a two-dimensional version of the freely rotating chain 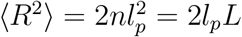.

Finally, we can also use the analytical results to describe the values of the tortuosity. By approximating the contour length at a given moment of time *t* as *v* · *t* and the end-to-end distance as the square root of the MSD given by Equation 8 with a correction factor of 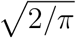, we get a good estimate of the tortuosity values (Figure 2c). This simple result predicts that for the short time scales of the ballistic and persistent regimes, the tortuosity is independent of growth speed and persistence and is essentially near the value of 1. At the times above the persistence time, the tortuosity scales as 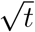, with the prefactor being independent of the growth velocity and inversely proportional to the square root of the persistence time or length (*L* ∝ *vt* and end-to-end distance 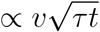), see Equation 15).

We should note that the success of the phenomenological run-and-tumble approach to quantify the growth trajectories is remarkable in the following sense. The simulated neurite growth is a rather complex process determined by non-trivial interactions of the forefront bead with ECM particles, that in turn depends on a number of other parameters, such as the link’s range, strength and life-time. One manifestation of these non-trivial interactions is the dependence (even if minor) of the observed growth speed on the sector angle. This example, however, indicates that certainly the run-and-tumble is a very useful, but still a bold approximation of a more complex neurite growth described here.

In an early rendition of this model, we were able to modify *θ* alone to describe the difference between growth patterns of neurites with a mutation in the MID1 gene and a wild type counterpart [19]. However, the other interaction parameters are also expected to contribute to the change in neurite growth statistics, and some complex cases might require the contribution of multiple parameters to describe the system. As such, we examined how modifications of these parameters influences growth, namely of the link formation range *R*, link persistence time *t*_link_ and strength *κ*_link_. The link formation range *R* mostly affects the average velocity, as increasing the range increases the chance of forming a long-range connection, thus effectively increasing the average link force (Equation 2). Therefore, increasing *R* correlates with an increase in *l*_*p*_, up to saturation (Supplementary Figure S3). Moving on to the link persistence time *t*_link_, a net increase in *l*_*p*_ is observed. This effect is likely observed because when *t*_link_ is lowered, the amount of links forming is increased, and so the average directionality of growth shifts closer to the average of 0 (Supplementary Figure S3). Finally, increasing *κ*_link_ shows an increase in *l*_*p*_ due to similar reasoning as with *R*. In addition to the growth cone parameters, the impact of the ECM parameters may also be studied. Increasing the radius of the ECM particles, for example, shows a decrease in MSD and an increase in tortuosity, up to saturation (Supplementary Figure S4). Other parameters such as the ECM area fraction will be explored in later sections.

In this subsection, the ECM particles are subjected to viscous forces and excluded volume interactions and can freely rearrange under the forces imposed by growing neurites. In this sense, such an ECM mimics a fluid-like viscoelastic medium. Additionally, however, we can also mimic viscoelastic solid-like matrices with our model.

### Effects of ECM geometry and stiffness

We next want to examine how the geometry of the extracellular space can impact the conformations of the growing neurites. To this end, we will consider ECM particles arranged in lattice-like configurations.

Furthermore, we “stabilize” the positions of the particles by connecting them by harmonic springs with equilibrium lengths equal to the distance between the particles *l*_lattice_, and a constant spring coefficient *κ*_lattice_. We start with the matrix arranged as a square lattice, in which each particle connected to its four nearest neighbors. In this solid-like scheme, the lattice geometry and parameters greatly influence growth trajectories (see supplementary video 2). We initially investigated the effects of *l*_lattice_ while keeping *κ*_lattice_ constant at 1.0, the sector angle at *θ* = 30°, and the rest of the parameters as was used for Figure 2 (see Table S1). We also set the ECM particle radius to *R*_ECM_ = 0.4, slightly smaller than the neurite radius of *R*_neurite_ = 0.5.

The homogeneous nature of this lattice allows us to better investigate how the ECM deforms due to neurite growth. The deformations were obtained by calculating the displacement of the particles from the first to the last frame, and averaging over 25 adjacent particles (Figure 3a). As the lattice spacing is defined from the center point of the ECM particles, we define a new effective lattice spacing 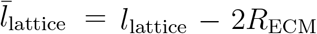, which describes the geometric spacing between each two ECM particles. The growth is quantified by the final values of the MSD and tortuosity and we plot them as a function of lattice density. Our results show that the MSD initially increases and then decreases as 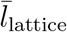 increases from 0.2 to 2.2 (Figure 3b). The final tortuosity shows the inverse relation, where it initially decreases and then increases (Figure 3c). The initial increase in the MSD (decrease in tortuosity) can be explained by the confining effect of the lattice. For 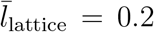, the free space between the lattice beads is smaller than the neurite diameter of 1.0, forcing the neurites to deform the lattice to advance. This, in turn, effectively increases the chances of strong turning events, due to the excluded volume interactions. As 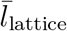 increases, the movement of the neurites through the lattice becomes less constricted. When the spacing is just enough to allow neurites to pass through, the ECM particles create a channeling effect, discouraging abrupt changes in directionality and thereby maximizing displacement. However, as spacing continues to increase, neurites gain more directional freedom, leading to increased directional changes and decreased global displacement. Thus, there exists an optimal lattice configuration that supports the most persistent growth.

**Figure 3.**
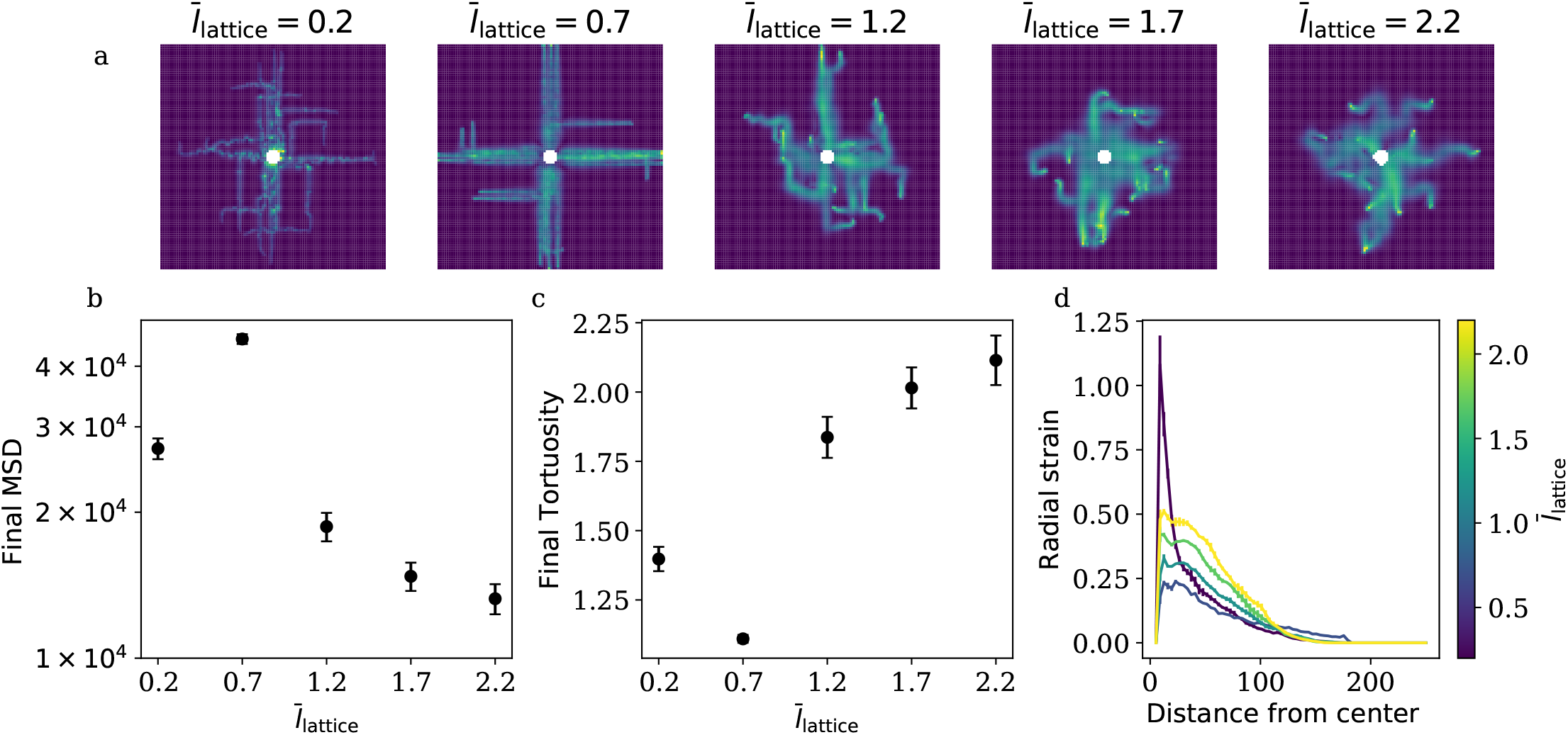
Square lattice simulations of neurite growth. Simulations were run for a total time of 40, with a timestep of 0.002, and an appropriate simulation box length of 500 to ensure all trajectories had sufficient simulation space to grow. Each condition included 78 averaged trajectories. Error bars represent standard error. Further details on parameters can be seen in Table S1. a) ECM deformation map calculated from the first and final frames of the run for each lattice spacing 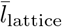. b-c) Final mean squared displacement and tortuosity over increasing lattice spacing. Both saturate at a lattice spacing of 1.5. d) Radially integrated ECM deformation, showing the maximum deformation intensity at each radial distance from the starting point of growth.

To further study this effect, we performed radial integration over the deformation maps, starting from the surface of the neuron cell body (Figure 3d). Deformation is maximal at 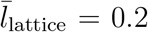, consistent with neurites having to displace the surrounding matrix to advance. At the ‘ideal’ configuration, ECM deformation is minimal, and the maps show only weak deformation along the neurite trajectories. Further increases in 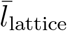 increase the average deformation, due to the increase in collision events resulting from the higher tortuosity.

In addition to 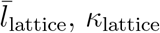 is also expected to influence lattice stiffness. Here, increasing *κ*_lattice_ shows the effect similar to increasing 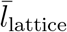, due to the net increase in lattice stiffness (see Supplementary Figure S5).

This model setting is similar to some recent work on polymer growth in mazes [28] and bacterial run-and-tumble movements in labyrinths, where thin channels not letting the swimming bacteria to reorient by tumbles are actually able to increase their persistence [29]. Other such ordered homogeneous lattices can also be studied in this model. In such cases, we expect to see similar results due to the similar homogeneity.

While such a regular type of an extracellular space can be realized in experimental setups, for example, in microfluidic systems or on micro-patterned surfaces, the most often used artificial ECMs are still disordered. Thus, we next will consider the case of random solid-like ECM and use this setting also to apply our model to explain the data on neurite growth in collagen gels of varying material properties.

### Stiffer ECMs support more persistent neurite growth in both *in vitro* experiments and simulations

Finally, we report on the experimental results investigating axonal growth in a 3D *in virto* setting and use these results to challenge our theoretical model.

In our experiments, primary hippocampal rat neurons were growing their neurites over a period of up to 4 hours, nested within layers of collagen hydrogels with varying concentration (see Figure 4a, Sup-plementary video 3 and Methods). Axonal growth cones were tracked at 2-minute intervals, resulting in one trajectory for each individual axon (see Figure 4b for summary or trajectories). Since the trajectories varied in the number of data points, we split them into equal-sized segments of 58 minutes. For each segment, the average instant velocity was determined, while the quotient of the end-to-end distance and the cumulative path length provided the tortuosity. We also define the related quantity of effective velocity as the end-to-end distance divided by the trajectory duration. Consequently, the average of the instant velocity, the tortuosity, and effective velocity were chosen as the parameters assigned to each axon (see Methods).

**Figure 4.**
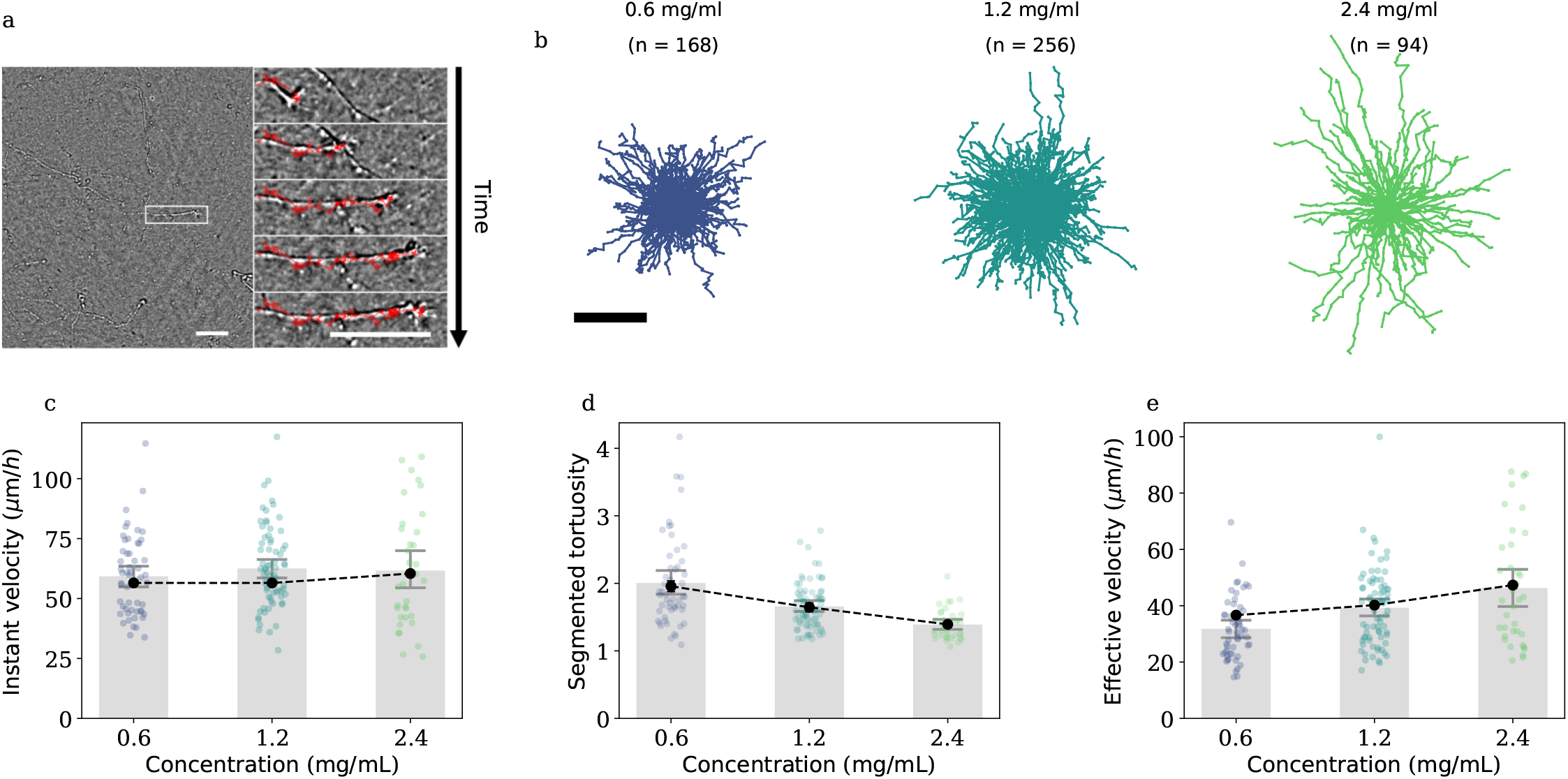
Simulation and experimental results of neurites growing in increasing collagen concentrations. a) Z-Projections of bright-field images (10x, 0.3 NA, z-stack height = 160 µm) of axons growing in a 3D collagen hydrogel (0.6 mg/ml). Highlighted area (white) is magnified (right), representing axonal growth over 160 minutes. Growth cone position was tracked every two minutes. Track is shown in red. Scale bar: 50 µm. b) 58-minute growth cone tracks at different collagen hydrogel densities (left: 0.6 mg/ml, n = 168; middle: 1.2 mg/ml, n = 256; right: 2.4 mg/ml, n = 94). Scale bar: 50 µm. c) Neurite instant velocity in experiments and simulation, as a function of collagen concentration. Colored dots represent experimental data, while the bar plot demonstrates the mean and standard error. Overlayed in black are the simulation results. d) Neurite tortuosity in experiments (colored dots) and simulations (black), as a function of collagen concentration. e) Effective velocity is defined as the end-to-end distance divided by the total trajectory duration. Conversion from collagen concentration in experiment to area fraction in simulations can be seen in Methods. Simulations averaged over 78 trajectories, with the parameters described in Table S1. Dashed lines included for visual guideline.

Our main focus was to study how the material properties of the gel affect the axonal growth. We observed that by increasing the concentration of the collagen (starting from 0.6 mg/mL and multiplied by a factor of 2) lead to an increase of the instant growth speed, but the effect was not too pronounced with an overall change of ~ 5% (see Figure 4b). In contrast, the persistence of growth (as quantified by tortuosity) was greatly affected (Figure 4c). Importantly, these results are in line with previous reports on a more persistent growth (lower tortuosity) on the surface of a more stiff substrate [1, 8], although this is now reproduced for the first time in a three-dimensional matrix. Here, the difference is of the order of 25%, much more significant than the increase in the velocity. The resulting increase in effective velocity corresponds to an increased outgrowth for neurites in stiffer environments. This resembles durotropic growth, as observed for pollen tubes with a solid or semisolid transmitting tract in plants [30].

To replicate this system using our simulation model, a Delaunay triangulation lattice ECM was employed, as illustrated in Figure 1d. Here, the dimensionless radius of the ECM particles *R*_ECM_ was chosen to be 0.25, in order to be able to implement potentially higher area fractions of ECM. We set the interaction parameters at the following constant values: *θ* = 30°, *R* = 20, *t*_link_ = 0.1, *κ*_link_ = 1.0. These parameters were chosen as they are known to reach similar persistances as the experimental setup (Figure 2c, Supplementary Figure S2) by the maximum simulation time when neurites reached the boundary of the simulation box (for the full parameter list, see Table S1).

The two key parameters in the model responsible for replicating the effects of the increasing collagen concentrations are the area fraction of the ECM particles *ϕ*_ECM_ and the spring coefficient *κ*_link_ determining the ECM stiffness. In collagen gels, the characteristic pore size scales as an inverse square root of the collagen concentration [31]. This effect can be captured by increasing the number density of the ECM particles in our two-dimensional setting. Here, the pore size has the same scaling with the number density [32].

Collagen gel stiffness is known to scale quadratically with collagen concentration [33]. Interestingly, in a two-dimensional system of beads connected by springs, increasing the number density of beads does not affect its stiffness. Higher density of beads along the direction of strain reduces the effective spring constant, while the increase of the number of springs in the orthogonal direction increases the effective spring constant in a compensatory fashion. Therefore, to account for the effect of concentration on stiffness we also linearly increase spring constant of ECM *κ*_lattice_ in our model (linear increase in 2D will be equivalent to quadratic increase in the 3D system).

From here, the simulation was sampled in the same manner as the experiments, where the position of the tip of the neurite was used for the tracking. To compare the two, however, both spatial and temporal dimensions in the model were re-normalized to match the actual experimental dimensions in the following way. We chose an experimental value of 0.8*µ*m for the diameter of the axonal shaft, which is consistent with known literature values for rat hippocampal neurons in the range 0.1 ~ 2.0 *µ*m [34, 35, 36, 37] and it sets our space scale. To normalize the time scale, we calculated the average cumulative track length over time for both the normalized simulation and the experiment. From here, the average track velocities were extracted, and from their comparison, the time normalization was obtained. This procedure was only performed for the highest concentration, and the same normalization parameters were used for all other conditions.

In comparing the simulation and experimental trajectories, the data sampling performed on the experimental data was repeated for the simulations. Next, the average instant velocity and tortuosities were calculated. Finally, we tuned the increasing number density of ECM particles and the linear increase in the ECM spring constant to match the experimental data for varying collagen concentrations. Given relatively large error bars in the experimental data, we used a manual tuning of the few parameters we have to achieve a quantitative match to the data. The area fractions used to match the collagen concentrations were (0.19, 0.38, 0.76) and the spring constants (0.25, 0.5, 1.0) (see Table S1).

We are able to reproduce a slowly increasing growth velocity with increasing collagen concentration (Figure 4b). Likewise, the resulting tortuosities in the model closely match those in the experiments (Figure 4c, red symbols). Finally, also the effective velocity increase is captured in simulations.

## 4 Conclusions and Discussions

Our theoretical model provides a mechanistic approach to describing the dependence of neurite growth on its surrounding extracellular matrix. It shows that with changing neurite-ECM interaction parameters, neurite growth patterns are altered consistently with a simple phenomenological random-walk theory. Here we focused on the statistics of individual independently growing neurons. Such a setting can be used, for example, to distinguish growth patterns associated with particular genetic mutations [19]. In the unconnected heterogeneous lattice, the absence of strong feedback from the lattice to the ECM particles resulted in growth patterns driven by adhesion with the growth cone dynamics being the primary determinant of neurite growth. Consequently, growth-cone-ECM interaction parameters were directly responsible for the persistence of growth. Experimentally, such configurations in simulations could represent artificial ECMs composed of gel beads [20], or to any lattice arrangement with structural elements with weak enough coupling allowing them to be easily deformed.

In contrast, the simulations with the square lattice configuration with elastic-like properties demonstrated that adhesion forces may be no longer the dominant factor in growth persistence. In particular, mechanical constraints imposed by the ECM’s geometry can play a critical role in neurite path-finding. While this idealized square lattice does not directly represent any specific *in vitro* system, its simplicity makes it well suited for isolating and studying the effects of structural constraints, which can later inform interpretations of more complex ECM configurations [28].

A heterogeneous network connected via Delaunay triangulation, represents a more realistic ECM similar to crosslinked polymer networks such as hydrogels. Such configurations, in both *in silico* and *in vivo* contexts, offer closer analogies to the heterogeneous environments encountered in brain context. In our experiments on rat neurons growing axons in artificial ECM made of collagen gel, we observed that higher collagen density was associated with greater growth persistence. These findings align with previous studies [38] and suggest a possible mechanical basis for how neurites navigate media of varying stiffness to reach target regions [10, 8]. While these works looked on the growth of axons on top of substrates with varying mechanical properties, here we study the axonal growth embedded in a 3D hydrogel. Under very natural assumptions on the model parameters, we could fully recapitulate these results. These involved assuming the increasing area fraction of the ECM particles and increasing stiffness of the ECM. Importantly, combining highly controlled experimental conditions with quantitative biophysical model, we can suggest that indeed the observed effects on the neurite growth are of purely physical origin. While the growth itself is an active process (involving traction force dipole) the resulting phenotype of the growth is a passive response/guidance of the ECM.

The simplicity of the present modeling framework makes it readily extendable to more complex mechanical environments encountered in biological systems. One potential application is the study of spine regeneration in zebrafish larvae, where axons traverse a wound lesion to reconnect with the opposite side in a microenvironment strongly regulated by extracellular matrix stiffness [39]. In such cases, additional chemical signaling processes could be incorporated into the model to better recapitulate observed regeneration patterns [40].

Another relevant system involves mechanosensing neurites, which alter their orientation in response to changes in environmental stiffness [10]. Simulations could be performed to isolate the effects of lattice stiffness changes arising solely from ECM geometry and material properties (i.e. passive biophysical contributions), and compare them to orientation shifts driven by mechanosensitive responses and to altered chemical signaling upon entering stiffer regions, to disentangle the biophysical from the biochemical contributions.

Finally, the framework can be extended beyond neuritogenesis to synaptogenesis, where simulated neurites form synapses and assemble into complete neuronal networks. Such an approach enables investigation of complex mechanisms, including how specific genetic mutations associated with epilepsy syndromes alter synaptic clustering patterns, from a biophysical standpoint [41].

## 5 Methods

### 5.1 Simulation setup

The simulation model was implemented using C++. All the governing equations are provided in the main text. Particle spatial interactions were computed using a hash map with cells of size *R*_cell_ = *R*_neurite_. The hash map limits particle interactions and searches interaction partners within the specified cell range, thereby considerably speeding up simulations. For the excluded volume interactions, the cell range was set to include up to the adjacent cells, resulting in a search algorithm that accounts for only nine cells. For establishing adhesion links, the cell range was set as *R/R*_cell_, where *R* is the growth cone adhesion range. The Delaunay triangulation used for the connected Delaunay lattice was implemented using the Delaunator library for C++ (https://github.com/delfrrr/delaunator-cpp). Several simulation parameters were kept constant across all simulations. The simulation timestep dt was set to 2 · 10^−3^ to en-sure stability. The neurite spring constant *κ*_neurite_ and the growth cone spring constant *κ*_cone_ were set to 2.5 and 0.01, respectively. This ensured that the neurite chain was much stiffer than the growth cone adhesion and that the growth cone was flexible enough to stretch towards its target. The growth rate parameter *α* was set to 10.0 to prevent neurites from getting stuck if they could not find an adhesion site, while not significantly influencing the growth dynamics. The neurite spring equilibrium length *l*_neurite_ was chosen as 1.2, ensuring that two neurites of radius *R*_neurite_ = 0.5 did not intersect in their equilibrium state, but also did not allow ECM beads to pass through the chain. Finally, the excluded volume parameter *ϵ* was set to 3.0 for all simulations. Lower values allowed adhesion forces to overwhelm the interactions, while higher values. Simulation reporting output is provided at specified equal intervals. By default, the number of sampled steps was set to 1000 to reduce the storage space required for each simulation run. The simulation outputs include snapshot files of all simulation particles for plotting purposes and a tracking file containing the position of the growth cones at each sampled time step. Plotting of the simulation snapshots was performed using Ovito [42], while analysis of the neurite tracking was conducted using Python. To reduce short-timescale fluctuations, we perform a preliminary rescaling of the recorded trajectories. The sampled time step *dt* is correspondingly rescaled to an effective value of 20 *dt*. Parameters of the fits to the simulation results for the mean squared displacement (by Equation (13)), velocity autocorrelation (by Equation (11)), and persistence time (by Equations (12, 14)) were obtained using the lmfit library [43].

### 5.2 Neuronal cell culture

Primary hippocampal neuronal cultures were prepared from postnatal day 0–2 (P0–P2) rat pups. Cell suspensions were generated from rat hippocampi as previously described [44]. Briefly, whole brains were rapidly removed and transferred into ice-cold hippocampal dissection buffer (HDB; 6 mg/mL glucose, 10 mM saccharose, 25 mM HEPES, and 5 mM NaHCO_3_ in HBSS without MgCl_2_ and CaCl_2_; Life Technologies, Darmstadt, Germany). Hippocampi were dissected, washed three times in HDB, and subjected to enzymatic dissociation in HBSS containing 0.01% DNase I, 0.1% dispase II, 0.01% papain (Roche, Mannheim, Germany), and 12.5 mM MgSO_4_. Tissue was processed using a brain-specific program on the gentleMACS Octo Dissociator (Miltenyi Biotec, Bergisch Gladbach, Germany), incubated at 37°C for 30 min, mechanically dissociated again, and centrifuged at 800 rpm for 4 min.

The pellet was triturated in serum-free Neurobasal-A medium supplemented with 2% B27, filtered through a 70 *µ*m nylon strainer. Subsequently, an equal volume of HDB containing 4% bovine serum albumin (BSA; Amresco, Solon, USA) was added to the filtrate, and the mixture was centrifuged at 1100 rpm for 8 min at room temperature to remove debris. To enrich neurons, cells were pre-plated on uncoated flasks for 1 h at 37°C (5% CO_2_), allowing glial cells to adhere. The neuronal supernatant was collected, centrifuged at 800 rpm for 8 min, and the final pellet was resuspended in complete neuronal medium (Neurobasal-A containing 2% B27, 0.5 mM L-glutamine, and 1% penicillin–streptomycin).

### 5.3 Preparation of cell culture embedded in collagen ECM

Our protocol for preparing collagen hydrogels was adapted for primary neurons from a protocol previously described in [21]. Powdered Neurobasal A medium (United States Biological, cat. N1020-02.10) was reconstituted and used at 1x and 10x, with 10x referring to the tenfold concentration of normal 1x working concentration. Collagen type 1 hydrogels were prepared from acid-dissolved rat tail (R) and bovine skin (G1) collagen (Matrix Bioscience). Both collagen types were mixed ice-cold at a mass ratio of 1:2 and were dissolved in H_2_O together with NaHCO_3_ and 10× Neurobasal A medium so that the final concentration of NaHCO_3_ was 27.4 *µ*M, the final Neurobasal A medium concentration was 1x, and the final collagen concentration was either 0.6 mg/ml, 1.2 mg/ml or 2.4 mg/ml. The mixture was then adjusted to pH 7.4 using 1 M NaOH. Optionally, the required number of neuronal cells for a final cell concentration of 3.3 × 10^6^ cells/ml hydrogel were suspended in 20 *µ*l of 1x Neurobasal A medium and added to the mixture. 10 *µ*l of the cell-free collagen solution was pipetted in a 15-well *µ*-slide (Ibidi, Gräfelfing, Germany) with a glass bottom, and polymerized for 90 min at 5% CO_2_ and 37 °C, resulting in a 800 *µ*m thick collagen hydrogel layer. Afterwards, 10 *µ*l of the cell-containing collagen solution was pipetted on top of the polymerized collagen layer and also polymerized for 90 min at 5% CO_2_ and 37 °C. 45 *µ*l of full growth medium (Neurobasal A + 2% B-27 + 0.5 L-glutamine + 1% Pen-Strep) was then added to each well. The *µ*-slide was incubated overnight. After 20h (day1), the medium was replaced with growth medium additionally supplemented with 6 nM Cytosine *β*-D-arabinofuranoside hydrochloride (Ara-C) to suppress proliferation of non-neuronal cell types such as glial cells.

### 5.4 Experimental setup

Cell cultures were imaged between days 3 and 5. At this stage, numerous axons were growing into the bottom layer of the collagen hydrogel. To ensure that the axons had a total length of at least 500 *µ*m, bright field image stacks between a height of 140 *µ*m and 300 *µ*m and a z-distance of 2 *µ*m were acquired with a ASI RAMM microscope equipped with a 10X 0.3 NA objective (Olympus, Japan). During imaging, the *µ*-slide was mounted in an incubation chamber (TOKAI HIT CO., Japan) set at 37°C and 5% CO_2_. The cells were imaged for up to 4 hours, with each field of view (FOV) being recorded every 2 minutes.

### 5.5 Data analysis

Maximum and minimum intensity projections in the z-direction of the same image stacks were computed and further preprocessed by high-pass filtering and contrast enhancement. Axonal growth cones were tracked manually using the open-source program “clickpoints” [45], which yielded one track per axon. Tracks shorter than 58 minutes (30 tracking points), e.g., due to the axons growing into or out of the z-stack range, were discarded from the analysis. Remaining tracks were subdivided into segments of 58 minutes, shorter residuals were discarded. Axons predominantly growing in z-direction were additionally discarded, as determined by a diagonal distance of a bounding box, defined by minimum and maximum *x* and *y* coordinates of the trajectory, of less than 5 *µ*m. For each segment, three parameters were calculated: instantaneous velocity was computed as the total trajectory contour divided by 58 minutes. Effective velocity was computed as the distance between the first and final tracking point divided by 58 minutes. The tortuosity was computed as the total trajectory contour divided by the distance between the first and final tracking point. For each track longer than 116 minutes, values for each 58-minute segment were averaged. Area fraction for the simulation was estimated by calculating the fraction of both types of collagen mixed in preparing the hydrogel. For example, at the lowest collagen concentration, the mixture contained 94*µL* of collagen out of a total volume of 500*µL*, from which we then estimate that collagen takes 19% of the total area in the final mixture in our simulations.

## 7 Acknowledgments

This work was funded by the German Research Foundation (DFG) project 460333672 CRC1540 EBM.

M.K. contributed in writing the simulation and the manuscript. L.B. contributed in conducting and analyzing the experimental results and in writing the manuscript. K.K. contributed in providing the neuron cell cultures and in writing the manuscript. F.F., M.K.2 and S.F. contributed in the planning of the work and in writing the manuscript. P.D. contributed in implementations of the simulation model. B.F. contributed in the experimental results and analysis, and in writing the paper. V.Z. supervised the work, contributed in devising the simulation and theoretical models, and contributed writing the manuscript.

## 8 Ethics approval statement

All animal experiments were approved by the local animal care and use committee (TS-3/2023) and conducted in accordance with the European Communities Council Directive and the German Animal Welfare Act, Directive 2010/63/EU, and §4 Abs. 1 and 3 of Tierschutzgesetz.

## 9 Data availability

Simulation code is available in https://github.com/matharkrv/neurite_framework. Raw experimental data is available in Zenodo (DOI: 10.5281/zenodo.18669424).

