## Supplementary Information text for "In silico neuritogenesis model underpins mechanical interactions with extracellular matrix as determinants of persistent axonal growth in stiffer microenvironments"

### Supplementary information: In silico twin model of neuritogenesis reveals the effects of extracellular matrix on neurite outgrowth

*Mathar Kravikass* [1,2 * #], *Lars Bischof* [3 *], *Kristina Karandasheva* [4], *Federica Furlanetto* [5], *Pritha Dolai* [1,2,6], *Sven Falk* [5], *Marisa Karow* [5], *Katja Kobow* [4], *Ben Fabry* [3], *Vasily Zaburdaev* [1,2 #]

[1] Department of Biology, Friedrich-Alexander-Universität Erlangen-Nürnberg, Erlangen, Germany.
[2] Max-Planck-Zentrum für Physik und Medizin, Erlangen, Germany.
[3] Department of Physics, Friedrich-Alexander-Universität Erlangen-Nürnberg, Erlangen, Germany.
[4] Department of Neuropathology, Universitätsklinikum Erlangen, Friedrich-Alexander-Universität Erlangen-Nürnberg, Erlangen, Germany.
[5] Institute of Biochemistry, Friedrich-Alexander-Universität Erlangen-Nürnberg, Erlangen, Germany.
[6] current address: National Institute of Technology Karnataka, Surathkal, Department of Physics, Mangalore, India.
Equal contribution (*)
Corresponding authors (#) Email address:



Keywords: *neuritogenesis, extracellular matrix, random walks, growth persistence*

| Name | Symbol | Figure 2 | Figure 3 | Figure 4 |
|---|---|---|---|---|
| Neuron radius | $R_{\text{neuron}}$ | 1.0 | 1.0 | 1.0 |
| Neurite radius | $R_{\text{neurite}}$ | 1.0 | 1.0 | 1.0 |
| Neurite mobility | $\mu_{\text{neuron}}$ | 1.0 | 1.0 | 1.0 |
| Neurite chain equilibrium length | $l_{\text{neurite}}$ | 1.2 | 1.2 | 1.2 |
| Neurite chain spring coefficient | $\kappa_{\text{neurite}}$ | 5.0 | 5.0 | 5.0 |
| Neurite cone spring coefficient | $\kappa_{\text{cone}}$ | 0.01 | 0.01 | 0.01 |
| Growth rate | $\alpha$ | 1.0 | 1.0 | 1.0 |
| ECM particle radius | $R_{\text{ECM}}$ | 0.25 | 0.4 | 0.25 |
| ECM particle mobility | $\mu_{\text{ECM}}$ | 2.0 | 1.25 | 2.0 |
| ECM area fraction | $\phi_{\text{ECM}}$ | 0.8 | - | $0.19, 0.38, 0.76$ |
| ECM lattice spacing | $l_{\text{lattice}}$ | - | 1.0 - 3.0 | - |
| ECM lattice spring coefficient | $\kappa_{\text{lattice}}$ | - | 1.0 | $0.25, 0.5, 1.0$ |
| Link spring coefficient | $\kappa_{\text{link}}$ | 1.0 | 1.0 | 1.0 |
| Link persistence time | $t_{\text{link}}$ | 0.01 | 0.01 | 0.01 |
| Sector angle | $\theta$ | $10 - 60$ | 30 | 30 |
| Range | $R$ | 20 | 20 | 20 |
| Simulation time step | $dt$ | 0.002 | 0.002 | 0.002 |
| Simulation box size | $L$ | 450 | 500 | 400 |
| Total simulation time | $T$ | 50 | 40 | 30 |

Table S1: Table of simulation parameters, as shown in each of the figures.

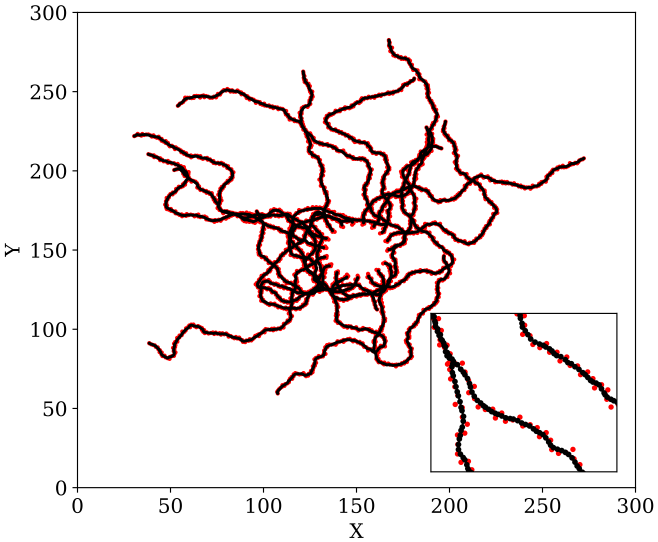


Figure S1: Comparison of individual neurite chain beads trajectories from center to the position of the front bead over time. Each solid line represents a different neurite in the simulation, and the points below represent the individual neurite chain beads.

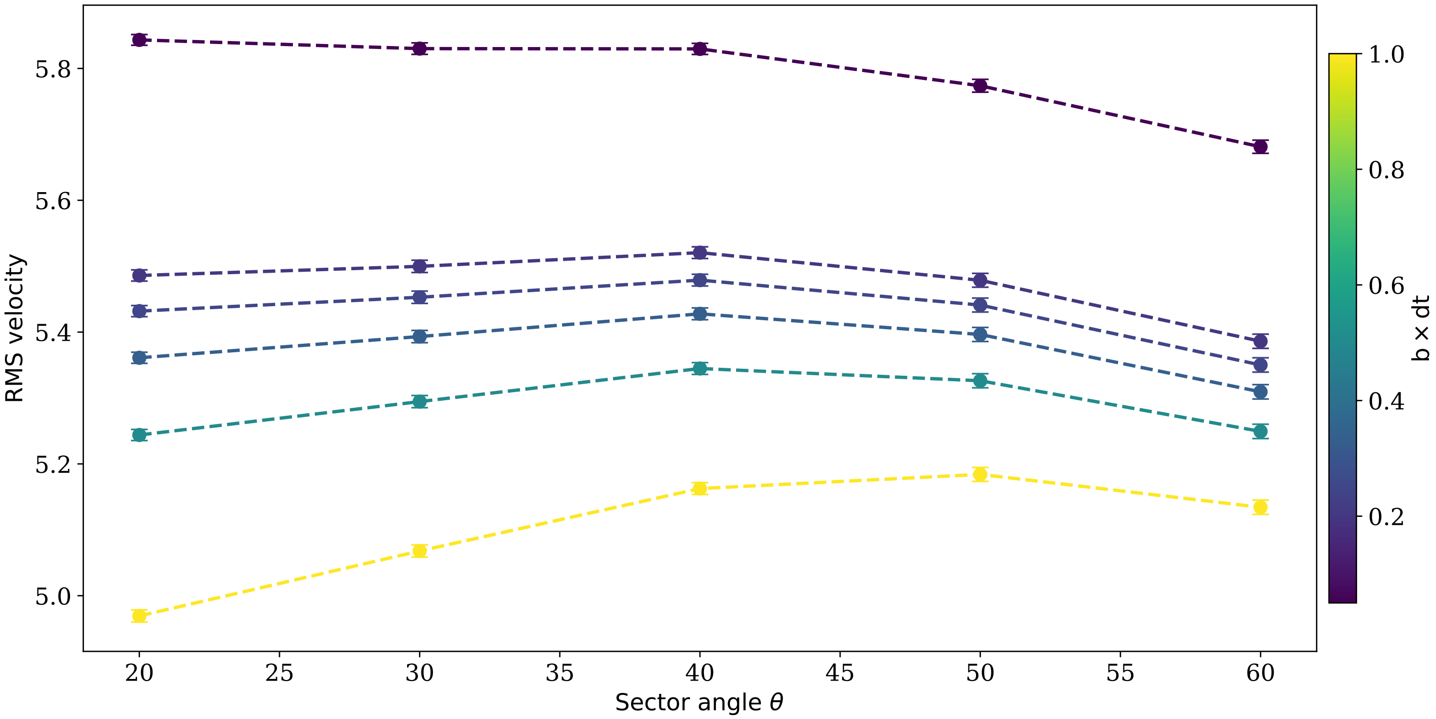


Figure S2: Root mean squared (RMS) velocity as a function of the sector angle $\theta$, using different time delays $b$. Parameters are as used for Figure 2 (refer to Table S1). Error bars represent the standard error. Increasing the time delay results with a decrease in the RMS velocities, with a slight shift in the curve. Regardless, the relative difference between the RMS velocities remains at less than 10%.

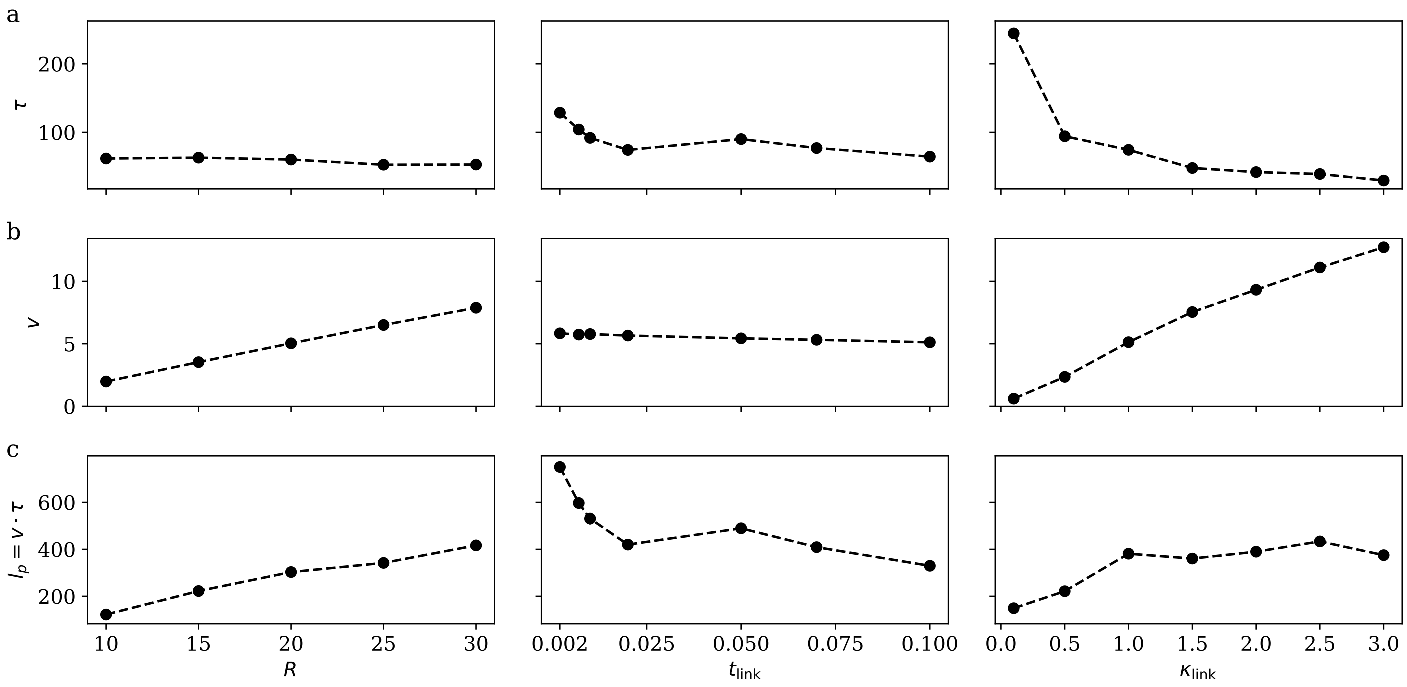


Figure S3: Simulation results averaged over 78 trajectories comparing the effect of the link range $R$, link coefficient $\kappa_{\text{link}}$ and link time $t_{\text{link}}$ on the model fitting. Parameters are as used for Figure 2 (refer to Table S1), with the exception of $R_{\text{ECM}} = 0.4$. a) Comparisons of the resulting $\tau$ persistence parameter from the model fitting. b) Comparisons of the calculated trajectory root mean squared velocity $v$ used for the model fitting. c) Comparisons of the persistence length $l_p$, obtained by $l_p = \tau \cdot v$. Dashed lines used for visualization purposes.

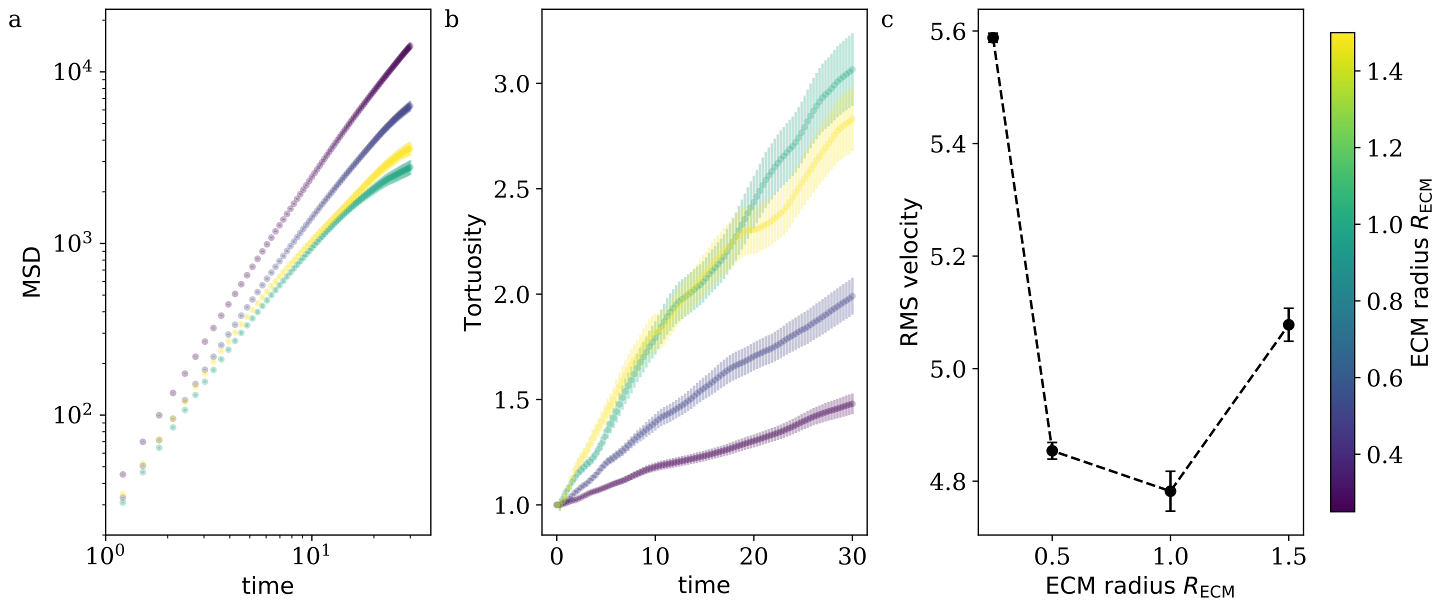


Figure S4: Simulations conducted with differing ECM radius $R_{\mathrm{ECM}}$, where the rest of the parameters are as described in Figure 2 (see also Table S1). Overall 78 trajectories were averaged for each condition. Error bars in all graphs represent the standard error. a) Mean squared displacement (MSD) as a function of time for different $R_{\mathrm{ECM}}$. Increasing $R_{\mathrm{ECM}}$ results in an decrease of MSD, which saturates at $R_{\mathrm{ECM}} = 1.0$. b) Tortuosity as a function of time for different $R_{\mathrm{ECM}}$. Compared to the MSD, the tortuosity increases with $R_{\mathrm{ECM}}$, with the same saturation point of $R_{\mathrm{ECM}} = 1.0$. c) Root mean squared (RMS) velocity as a function of $R_{\mathrm{ECM}}$. RMS velocity starts with an initial decrease, which then shifts to a weaker increase from $R_{\mathrm{ECM}} = 1.0$ onward. Dashed lines used for visualization purposes.

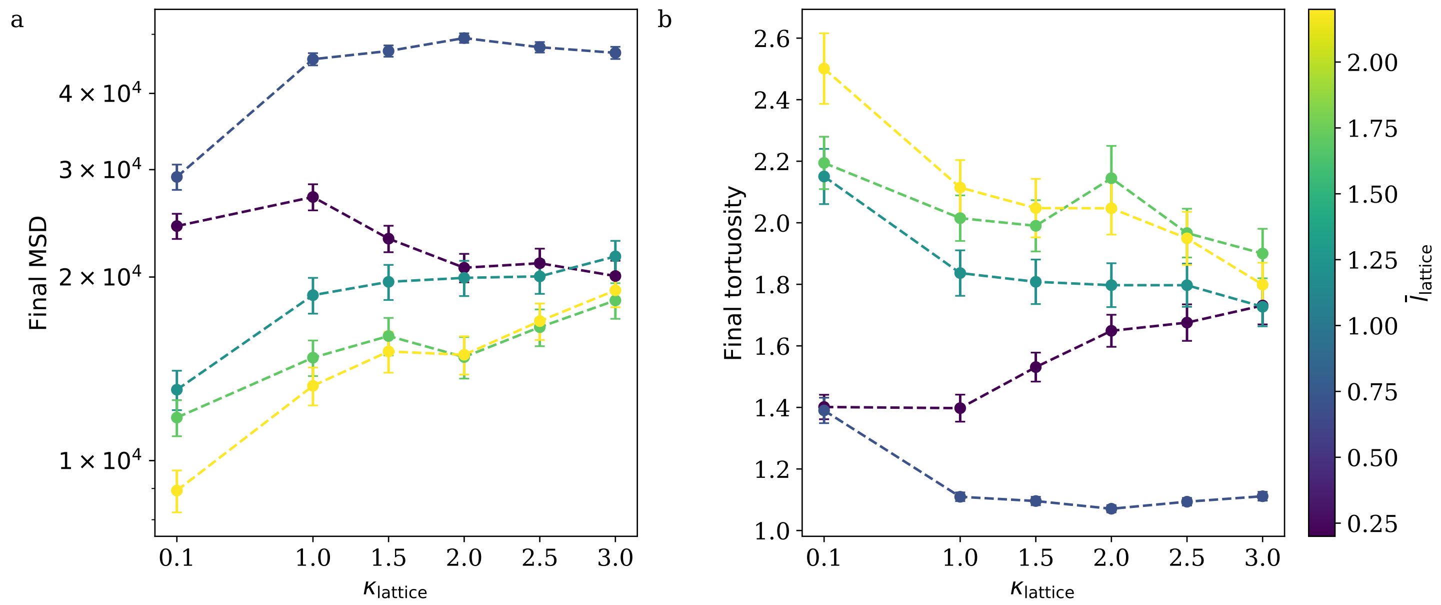


Figure S5: Comparisons of differing square lattice parameters averaged over 78 trajectories per condition, supplementary to Figure 3. Parameters are as used for Figure 3 (refer to Table S1). a) Final mean squared displacement (MSD) as a function of lattice spring constant $\kappa_{\mathrm{lattice}}$, for each simulated lattice spacing $\bar{l}_{\mathrm{lattice}} = l_{\mathrm{lattice}} - 2R_{\mathrm{ECM}}$. Except for $\bar{l}_{\mathrm{lattice}} = 1.0$, where growth is stunted due to the spacing being smaller than the neurite diameter, all other conditions exhibit an increase in MSD with $\kappa_{\mathrm{link}}$. b) Similarly to the MSD, in exception for the case of $\bar{l}_{\mathrm{lattice}} = 1.0$ all tortuosities decrease with $\kappa_{\mathrm{link}}$, indicating a more persistent growth is accompanied with the increased stiffness.
